# Microplastic shape, concentration and polymer type affect soil properties and plant biomass

**DOI:** 10.1101/2020.07.27.223768

**Authors:** Yudi M. Lozano, Timon Lehnert, Lydia T. Linck, Anika Lehmann, Matthias C. Rillig

## Abstract

Microplastics are an increasing concern in terrestrial systems. These particles can be incorporated into the soil in a wide range of shapes and polymers, reflecting the fact that manufacturers produce plastics in a variety of physical and chemical properties matching their intended use.Despite of this, little is known about the effects that the addition into the soil of microplastics of different shapes, polymer type and concentration levels may have on soil properties and plant performance.

To fill this gap, we selected four microplastic shapes: fibers, films, foams and fragments; and for each shape we selected three microplastics made of one of the following polymers: polyester, polyamide, polypropylene, polyethylene, polyethylenterephthalat, polyurethane, polystyrene and polycarbonate. In a glasshouse experiment, each microplastic was added to a soil from a dry grassland at a concentration of 0.1%, 0.2%, 0.3% and 0,4%. A carrot (*Daucus carota*) plant grew in each pot during four weeks. At harvest, shoot and root mass, soil aggregation and microbial activity were measured.

Our results showed that all microplastic shapes increased shoot and root masses. As concentration increased, microfibers increased plant biomass probably as fibers may hold water in the soil for longer. In contrast, microfilms decreased biomass with concentration, likely because they can create channels in the soil that promote water evaporation affecting plant performance. All microplastic shapes decreased soil aggregation, probably since microplastics may introduce fracture points in the aggregates affecting their stability and also due to potential negative effects on soil biota. The latter may also explain the decrease in microbial activity with, for example, polyethylene films. Similar to plant biomass, microfilms decreased soil aggregation with increasing concentration.

Our study tested the microplastic shape mediation and dissimilarity hypotheses, highlighting the importance of microplastic shape, polymer type and concentration when studying the effects of microplastics on terrestrial systems.

## INTRODUCTION

Microplastics (< 5 mm) are increasingly reported in terrestrial systems, and due to slow turnover, may be gradually increasing through additions including soil amendments, plastic mulching, irrigation, flooding, atmospheric input and littering or street runoff ^1-4^. As a result of their manufacturing origin, environmental degradation and intended use, these particles occur in many shapes, and cover a high physical and chemical diversity ^5, 6^.

Each microplastic shape may be represented by different polymer types as manufacturers seek to produce plastic with specific properties (e.g., flexibility, roughness, resistance, durability) ^7^. However, these polymer types are composed of different monomers, which can potentially be hazardous for the environment ^8^. For instance, polyurethane, a polymer used to produce flexible foams, is made of monomers highly toxic for humans ^8^ and potentially for soil biota as millions of tons of this plastic are produced annually, potentially increasing its concentration in the soil.

Agricultural soils are particularly prone to being enriched with microplastic, as several pathways for plastic addition and incorporation exist in agroecosystems. For example, fibers are found in soil amended with sewage sludge ^9^. Indeed, microplastic concentrations of 30.7×10^3^ particles kg^-1^ dry sludge have been reported ^10^. Similarly, plastic mulching is widely used in certain types of agricultural fields ^2, 11^, and thus microplastic film concentrations in soil may increase ^11^. The wide-spread application and the intentional or unintentional ubiquitous distribution of plastics affects even remote landscapes and agricultural sites with plastic-free management plans ^12^. Other microplastic shapes, such as foams or fragments can be incorporated into the soil due to littering, street runoff ^2^ or wind deposition ^3, 13^.

Our knowledge about microplastic effects on terrestrial systems is still scarce and effects reported for plants and soil often seem contradictory, as effects may differ depending on microplastic shape, polymer structure, degradation, additives and concentration, as well as on the target plant or soil. For instance, microplastic granules of ethylene propylene at 5% may decrease plant biomass likely linked to its polymer composition ^14^, while polyester fibers at 0.4% may have the opposite effect as microfibers can enhance soil water content and soil aeration ^15^. Polyester fibers can also increase soil aggregation as they may help to entangle soil particles ^15, 16^, while opposite effects are detectable for e.g. polyamide fibers ^15^. Likewise, microplastics can affect soil microbial activity as they can increase mortality and histological damage in soil macroorganisms ^17, 18^, and decline richness and diversity of bacterial communities as seen with polyethylene films ^19, 20^.

As microplastics differ in a number of properties, including shape, polymer type and concentration, effects on plant species and soils may differ as a function of these properties. To test this, we established a glasshouse experiment that included four microplastic shapes (i.e., fibers, films, foams and fragments), each of them with three different polymer types and four concentrations (0.1, 0.2, 0.3, 0.4% w/w). We evaluated effects on shoot and root masses of the plant *Daucus carota*, and on soil aggregation and soil microbial activity. In doing so, we tested the shape dissimilarity and shape mediation hypotheses proposed by Rillig et al ^5^ : effects are predicted to be strongest when microplastic particles diverge most clearly from the population of naturals shapes, and at equity of shape effects will be mediated by physical/ chemical properties of the particles.

## MATERIALS AND METHODS

### Species selection

As microplastics could affect soil water status ^15^, we selected *Daucus carota* (wild carrot), a biennial herbaceous typical from dryland ecosystems ^21^ that exhibits clear responses to water availability ^22^. Seeds of this plant species were obtained from a commercial supplier in the region (Rieger-Hofmann GmbH, Blaufelden, Germany).

### Microplastics

We selected twelve different secondary microplastics representing four microplastic shapes: fibers, films, foams and fragments, and eight polymer types: polyester (PES), polyamide (PA), polypropylene (PP), polyethylene (PE), polyethylenterephthalat (PET), polyurethane (PU), polystyrene (PS) and polycarbonate (PC).

Within each microplastic shape we selected three microplastics made of different polymers. That is, as fibers we selected: Polyester (PES, Rope Paraloc Mamutec polyester white, item number, 8442172, Hornbach.de), polyamide (PA, connex, item number 10010166, Hornbach.de) and polypropylene (PP, Rope Paraloc Mamutec polypropylene orange, item number, 8442182, Hornbach.de); as films we selected: Polyethylene low-density (PE, silo film black, folien-bernhardt.de), Polyethylenterephthalat (PET, company: Toppits / product: oven bag), and Cast Polypropylene (PP, company: STYLEX / product: transparent folders); as foams we selected: Polyurethane (PU, grey foam sheet, item number, 3838930, Hornbach.de), polyethylene (PE, black low density closed cell ETHAFOAM polyethylene foam, Rugen, shrugen.en.alibaba.com), and expandable Polystyrene (PS, EPS70 Insulation Packing Board SLABS, Wellpack Europe). Finally, as fragments we selected: Polyethylenterephthalat (PET, VioStill, item number 41005958, vio.de), Polypropylene (PP, black plastic pots, treppens.de) and Polycarbonate (PC, CD-R Verbatim).

Fibers and films were manually cut with scissors. A length of 5.0 mm or 5.0 mm^2^, respectively, were established as an upper size threshold in order to generate microplastic fibers and films. Foams and large solid plastics were cut into small pieces by using a Philips HR3655/00 Standmixer (1400 Watt, ProBlend 6 3D Technologie, Netherlands), sieved through 4 mm mesh and, if necessary, cut with scissors in order the obtain microplastic foams and fragments (i.e., <5mm^2^). Microplastics were microwaved (2 min at 500 watts) to minimize microbial contamination. Temperature did not approach melting points during microwaving.

### Soil preparation

We collected a sandy loam soil (0.07% N, 0.77% C, pH 6.66) from a dry grassland located in Dedelow, Brandenburg, Germany (53° 37’ N, 13° 77’ W). The soil was air-dried, sieved (4 mm mesh size), homogenized and then mixed with each of the microplastics at a concentration of 0.1, 0.2, 0.3 and 0.4% (w/w). Thus, 0.19, 0.38, 0.57 and 0.76 g of each microplastic type were mixed into 190 g of soil for each pot (4 cm diameter, 21 cm height, 200 ml). Soil preparation was done separately for each pot. Microplastics were separated manually and mixed with the soil during 1 min in a large container, before placing it into each individual pot, to help provide an equal distribution of microplastics throughout the soil. Soil was mixed in all experimental units, including the controls, for the same amount of time and with the same intensity, in order to provide the same disturbance.

### Experimental design

In October 2019 we established the experiment in a glasshouse with a daylight period set at 12 h, 50 klx, and a temperature regime at 22/18 °C day/night with a relative humidity of ∼40 %. Prior to seedling transplanting, pots were incubated for two weeks allowing the interaction between the soil microbial communities and the microplastic particles as well as the potential leaching of plastic components into the soil. During that time, pots were well-watered twice a week by gently spraying 50 ml of distilled water onto the soil surface. Seeds of *Daucus carota* (∼ 1000 seeds) were surface-sterilized with 4 % sodium hypochlorite for 5 min and 75% ethanol for 2 minutes and then thoroughly rinsed with sterile water. The seeds were germinated in trays with sterile sand, and individual seedlings of similar size were transplanted into pots three days after germination. One seedling was added per pot. Following, pots were watered for 4 additional weeks. We thus had 12 microplastic types (i.e., 4 microplastic shapes x 3 polymer types) x 4 concentration levels x 7 replicates = 336 pots. Fourteen additional pots were established as a control without microplastics. All pots were randomly distributed in the greenhouse chamber, and their position shifted twice during the experiment to homogenize environmental conditions. At harvest, plants were separated into above and belowground parts; soil was divided into two subsamples of ∼30 g each, one was air-dried and stored at ∼25 °C for soil aggregation analyses and the other was kept at 4 °C for a maximum of 1 month for soil microbial activity analyses.

### Measurements

#### Biomass

Roots were carefully removed from the soil and gently washed by hand. Then, shoots and roots were dried at 60 °C for 72 h, after which their mass was determined.

#### Soil aggregation

Water stable soil aggregates as a proxy of soil aggregation were measured following a protocol by Kemper and Rosenau ^23^, modified as described in Lehmann et al ^24^. Briefly, 4.0 g of dried soil (<4 mm) was placed on small sieves with a mesh size of 250 μm. Soil was rewetted with deionized water by capillarity and inserted into a sieving machine (Agrisearch Equipment, Eijkelkamp, Giesbeek, Netherlands) for 3 min. Agitation and re- wetting causes the treated aggregates to slake. The water-stable fraction (dry matter) was dried and weighed. Subsequently, we extracted the coarse matter which also was dried at 60 °C for 24 h. Soil aggregation (i.e., water stable aggregates) was calculated as: WSA (%) = (Dry matter- coarse matter)/(4.0 g - coarse matter).

#### Soil microbial activity

We measured soil respiration as it is considered a good proxy of total microbial activity ^25^. MicroResp™, as described by Campbell et al ^26^, was used to measure community respiration. To do so, we placed approximately 0.42 g of soil into each well of the 96-deep well plates. Four wells were used for each treatment (technical replicates). Soil samples were incubated for 1 day at 25 °C prior to carrying out the assay. CO_2_ detection plates were read and then the deep-well plates were sealed with the pre-read CO_2_ detection plates and incubated at 25 °C for 6 h in the dark, as recommended by the manufacturer (Macaulay Scientific Consulting, UK). The change in absorbance values after incubation was then measured on a spectrophotometer microplate reader (Benchmark Plus Microplate Spectrophotometer System, BioRad Laboratories, Hercules, CA, US) at a wavelength of 570 nm. The CO_2_ rate (µg CO_2_-C g^-1^ h^-1^) per well was calculated using the formula provided in the MicroResp™ manual (Macaulay Scientific Consulting, UK).

### Statistical analyses

The effect of microplastic shape, polymer type and concentration on shoot and root masses, soil aggregation and microbial activity were explored through linear models and multiple comparisons “multcomp package”. First, residuals of the linear models were checked to validate assumptions of normality and homogeneity. When necessary, we implemented the function “varIdent” to account for heterogeneity in the treatment. Then, to the selected model, we implemented the function “glht” and the “Dunnett” test from the “multcomp” package ^27, 28^, in order to compare each microplastic treatment with the control (without microplastics). Additionally, effect sizes were estimated to show the variability in the response of our variables, by comparing each microplastic type (i.e., shape and polymer) with the control pots (without microplastics) for each concentration level, using a bootstrap-coupled estimation “dabestr package’ ^29^.

Finally, we tested the shape dissimilarity hypothesis by determining the effects of the same polymer type from different shapes on our response variables. To do so, we chose the treatments where polypropylene, polyethylenterephthalat and polyethylene as different microplastic shapes were available for these polymers. We performed an analysis of variance “aov” that included shape and polymer as fixed factor. Shoot mass was log-transformed to meet normality assumptions. Then, we performed multiple comparisons “glht” among treatments by using the “Tukey” test and the function “sandwich” from the eponymous package; this function provided a heteroscedasticity-consistent estimate of the covariance matrix ^28, 30^. Statistical analyses were done in R 3.5.3 ^31^.

## RESULTS

### Shoot mass

Shoot mass was affected by microplastic shape, polymer type and concentration. Films, fragments, foams and fibers increased shoot mass in that order (Figure 1a, Table S1). This was also true for all polymer types except polyamide and polystyrene whose effects were similar to control (Figure 1a, Table S2). Fibers increased shoot mass with increasing concentration (Figure 1b), a pattern mainly observed with fibers made of polypropylene (PP) (Figure 1b, Table S2). Microplastic films overall, increased shoot mass (Figure 1a, Table S1). However, the trend was contrary compared to fibers (Figure 1b): the lower the concentration of microplastic films the more positive the effect shown for polypropylene (PP) and polyethylenterephthalat (PET) films (Figure 1b, Table S2). Microplastic foams had contrasting effects depending on the polymer type. That is, polyethylene (PE) and polystyrene (PS) tended to decrease shoot mass with increasing concentration while polyurethane (PU) showed no obvious pattern (Figure 1c, Table S2, S3). Although microplastic fragments overall increased shoot mass no clear concentration pattern was present (Figure 1c, Table S1).

**Figure 1.**
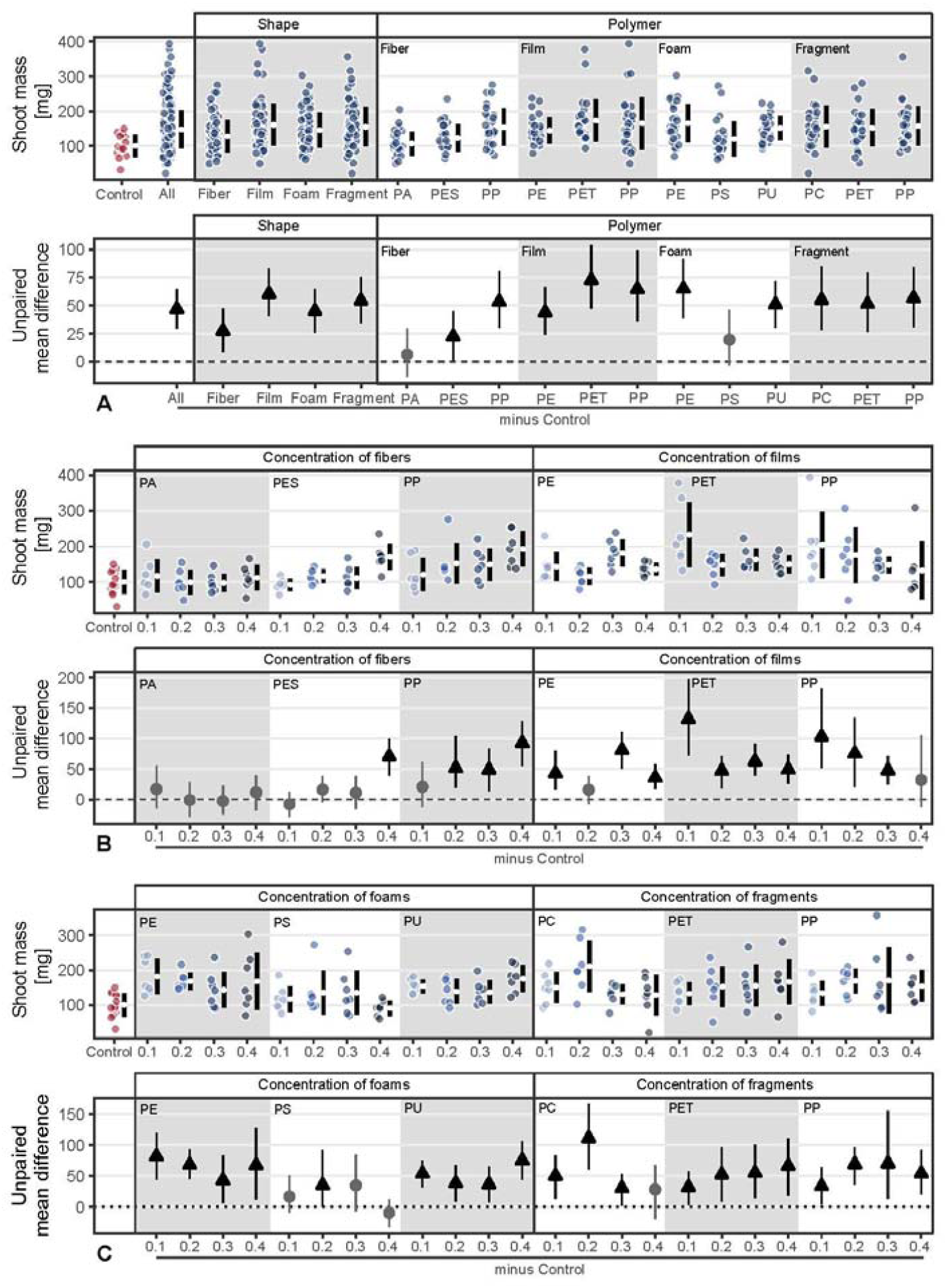
Shoot mass response to microplastic shape and polymer type at different concentrations (0.1%, 0.2%, 0.3% and 0.4%). Effect sizes and their variance are displayed as means and 95% confidence intervals. Effects are color-coded: gray circles indicate neutral effect sizes, black arrows with an arrow head pointing upwards indicate positive effects; no negative effects were detected. Horizontal dotted line indicates the mean value of the control. Polymers : PA (polyamide), PES (polyester), PP (polypropylene), PE (polyethylene), PET (Polyethylenterephthalat), PS (polystyrene), PU (polyurethane) and PC (polycarbonate). See statistical comparisons in Table S1-S3; n = 7 for microplastics, n=14 for control samples.

### Root mass

Root mass was affected by microplastic shape, polymer type and concentration. Films, foams and fragments increased root biomass while fibers led to a similar biomass to the control (Figure 2a, Table S1). Although root mass was positively affected by microplastics, the effects diverged from those found for shoot mass. Root mass was only altered by the addition of microfibers under the highest concentration, especially for polyamide (PA) fibers, causing a positive impact (Figure 2b, Table S3). Microplastic films and fragments increased root mass irrespective of the concentration (Figure 2b), while for foams we found an increase in root mass as concentration increased for polyurethane (PU); root mass tended to decrease for polyethylene (PE) and polystyrene (PS) (Figure 2c, Table S3).

**Figure 2.**
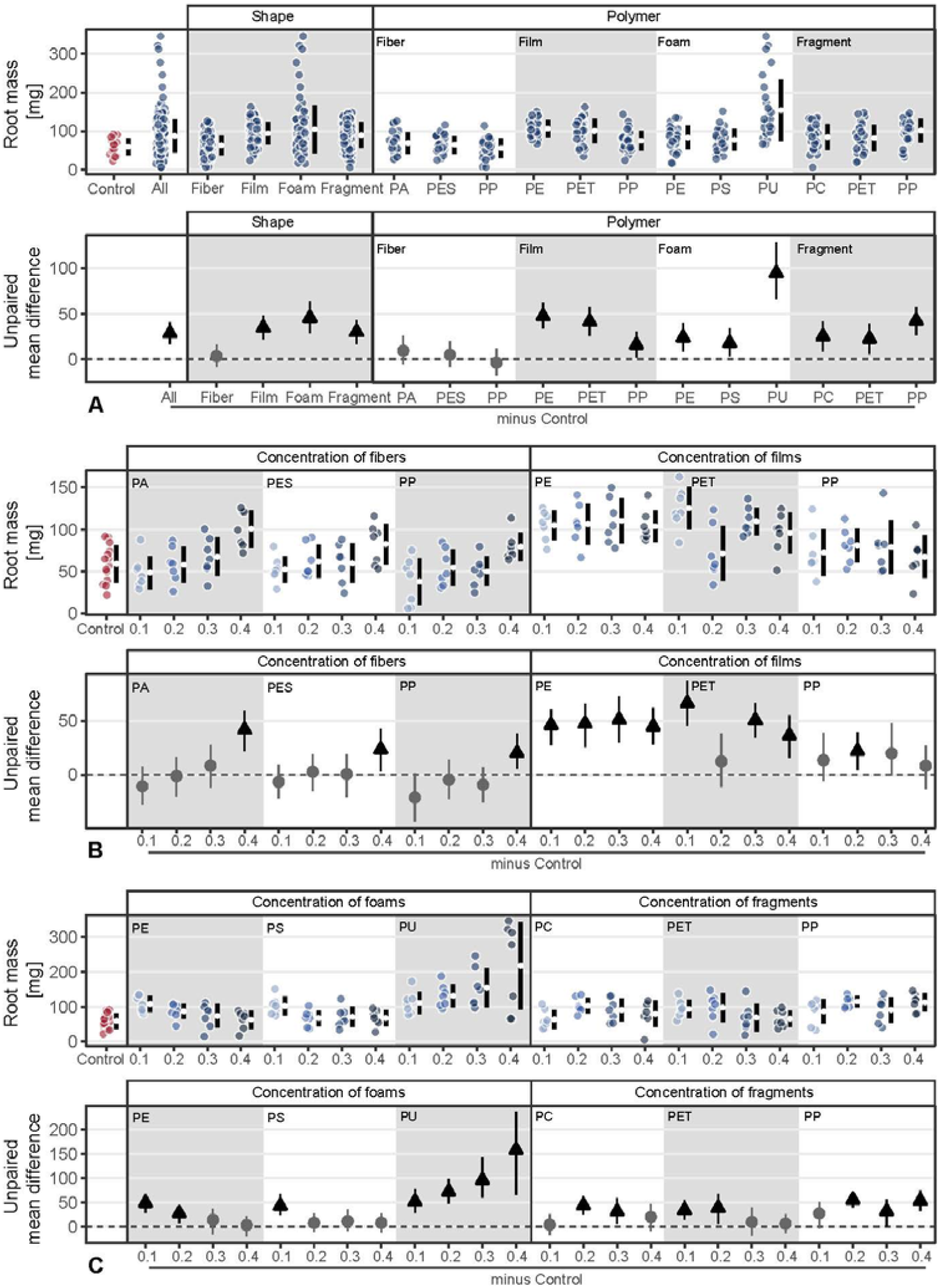
Root mass response to microplastic shape and polymer type at different concentrations (0.1%, 0.2%, 0.3% and 0.4%). Effect sizes and their variance are displayed as means and 95% confidence intervals. Effects are color-coded: gray circles indicate neutral effect sizes, black arrows with an arrow head pointing upwards indicate positive effects; no negative effects were detected. Horizontal dotted line indicates the mean value of the control. Polymers: PA (polyamide), PES (polyester), PP (polypropylene), PE (polyethylene), PET (Polyethylenterephthalat), PS (polystyrene), PU (polyurethane) and PC (polycarbonate). See statistical comparisons in Table S1-S3; n = 7 for microplastics, n=14 for control samples.

### Soil aggregation

Overall, the different microplastic shapes and polymer types negatively influenced soil aggregation (Figure 3a). Microplastic fibers consistently reduced the stability of soil aggregates, irrespective of concentration and polymer. For microplastic films and foams, we found concentration dependent trends: microplastic films reduced soil aggregate stability while foams, especially polyethylene (PE), had the opposite pattern as concentration increased (Figure 3b,c). Microplastic fragments showed no clear pattern with concentration: polyethylenterephthalat (PET) and polypropylene (PP) fragments reduced soil aggregation at lower concentrations but at the highest concentration (0.4%) this effect was neutralized (Figure 3c). Polycarbonate (PC) fragments showed a non-linear concentration effect, with the highest concentration causing strong reduction in soil aggregate stability (Figure 3c). Similar to shoot mass, soil aggregation increased with fibers but decreased with films as concentration increased (Figure 3b).

**Figure 3.**
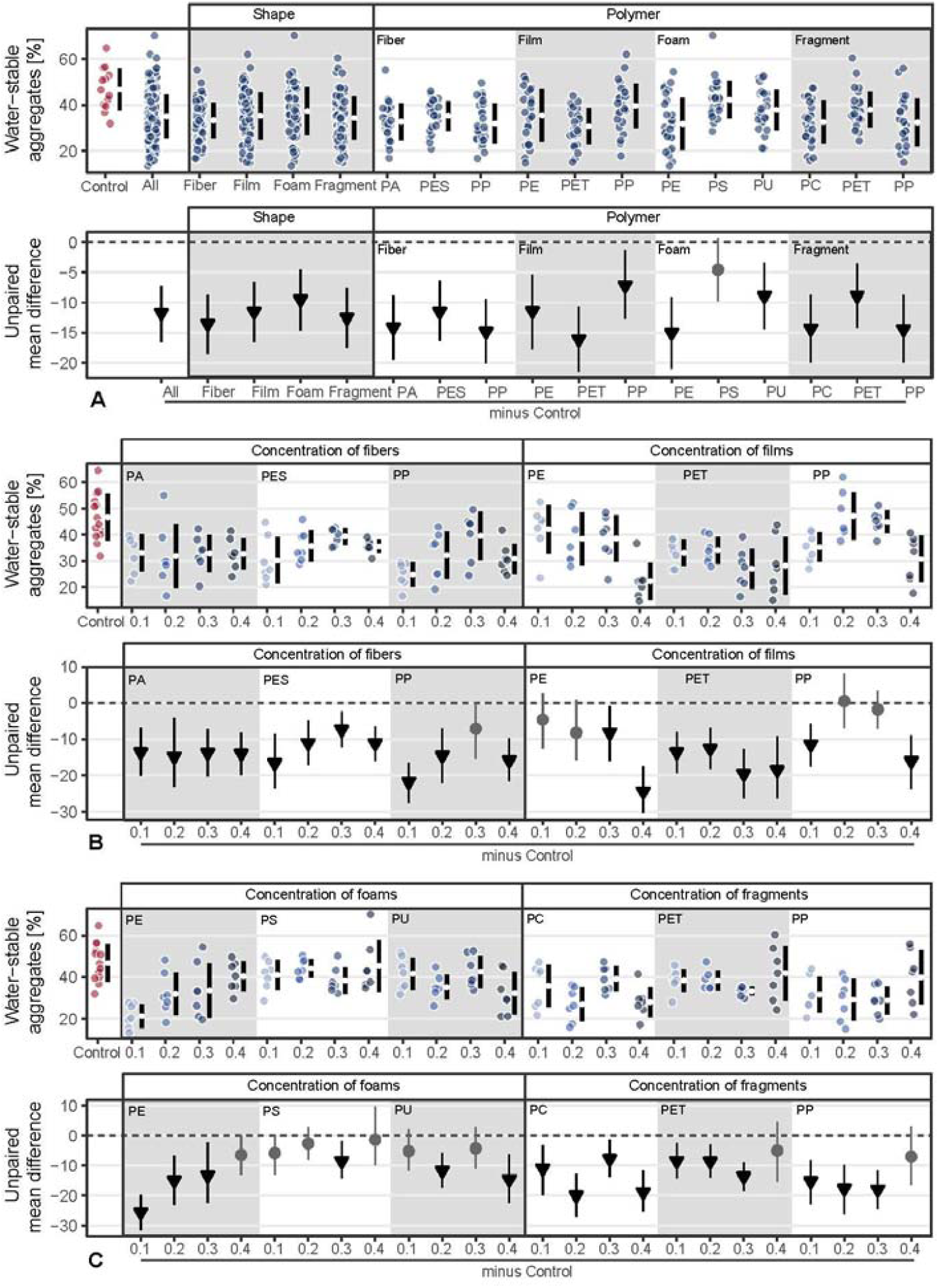
Soil aggregation (ie., water stable aggregates) response to microplastic shape and polymer type at different concentrations (0.1%, 0.2%, 0.3% and 0.4%). Effect sizes and their variance are displayed as means and 95% confidence intervals. Effects are color-coded: gray circles indicate neutral effect sizes, black arrows with an arrow head pointing downwards indicate negative effects; no positive effects were detected. Horizontal dotted line indicates the mean value of the control. Polymers: PA (polyamide), PES (polyester), PP (polypropylene), PE (polyethylene), PET (Polyethylenterephthalat), PS (polystyrene), PU (polyurethane) and PC (polycarbonate). See statistical comparisons in Table S1-S3; n = 7 for microplastics, n=14 for control samples.

### Microbial activity

The effect of microplastics on microbial activity was highly variable (Figure 4, Table S1). We observed that polyethylene (PE) films and polypropylene (PP) fragments decreased microbial activity (Figure 4a). Regarding concentrations, fibers decreased microbial activity at higher concentrations, i.e., at 0.3% for polyamide (PA) and polyester (PES) and at 0.4% for polypropylene (PP). By contrast, low concentrations of PA fibers, i.e., 0.1%, increased microbial activity. Microplastic films and foams had an overall neutral or negative effect on microbial activity, respectively (Figure 4a). Only polyethylenterephthalat (PET) films at 0.2% concentration and foams made of polyurethane (PU) at 0.2%, polyethylene (PE) at 0.3% and polystyrene (PS) at 0.4% had a positive effect on microbial activity. Overall, microplastic fragments had a neutral or negative effect but polycarbonate (PC) and PET at intermediate values had a positive effect on microbial activity (Figure 4d).

**Figure 4.**
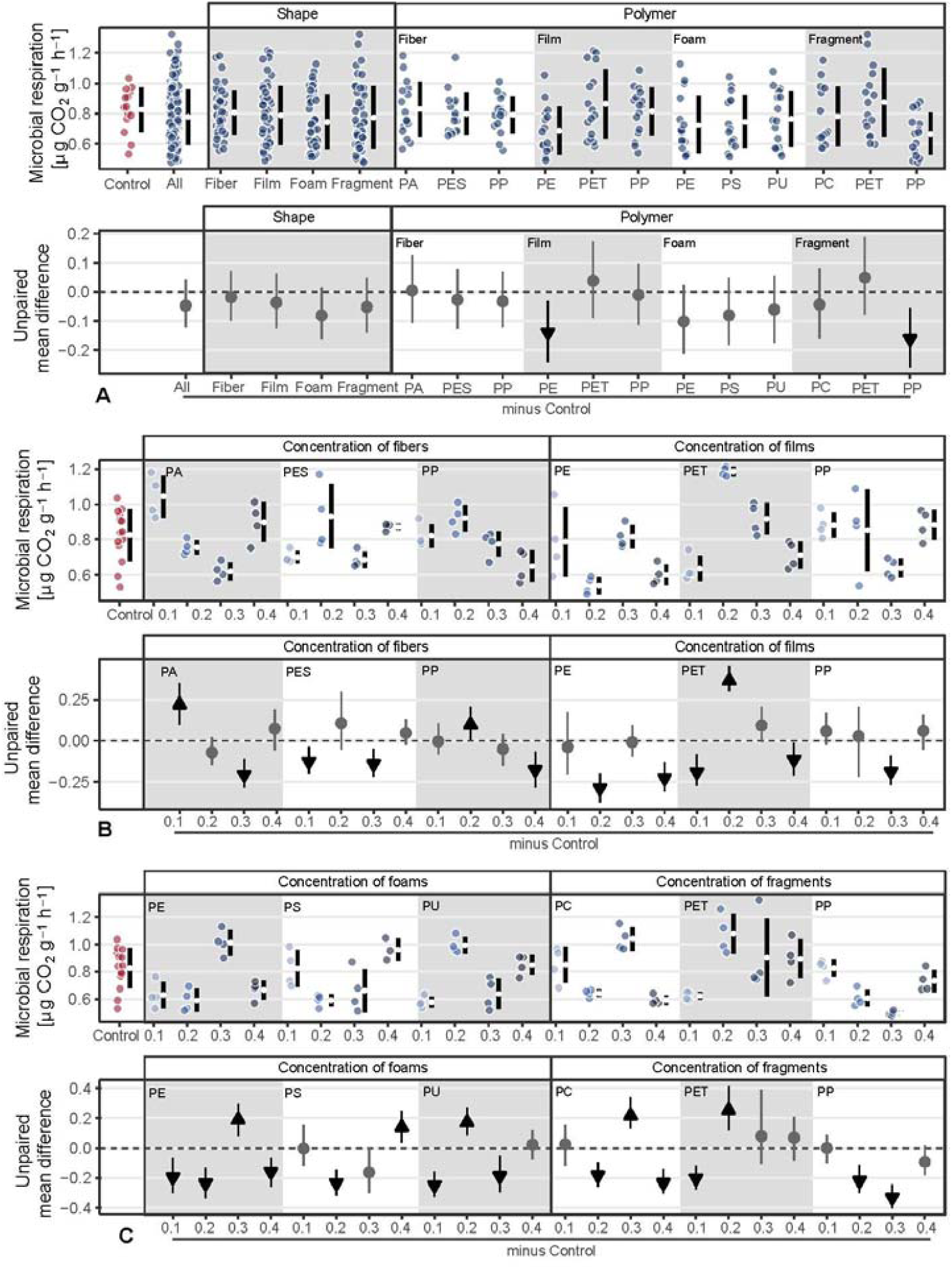
Microbial activity (ie., microbial respiration) response to microplastic shape and polymer type at different concentrations (0.1%, 0.2%, 0.3% and 0.4%). Effect sizes and their variance are displayed as means and 95% confidence intervals. Effects are color-coded: gray circles indicate neutral effect sizes, black arrows with an arrow head pointing upwards or downwards indicate positive or negative effects, respectively. Horizontal dotted line indicates the mean value of the control. Polymers: PA (polyamide), PES (polyester), PP (polypropylene), PE (polyethylene), PET (Polyethylenterephthalat), PS (polystyrene), PU (polyurethane) and PC (polycarbonate). See statistical comparisons in Table S1-S3; n = 7 for microplastics, n=14 for control samples.

### Microplastics shape-related hypothesis

Finally, in addition to the shape mediation hypothesis, we tested the shape dissimilarity hypothesis. Our results showed the importance of microplastic shape over polymer type for different plant traits and soil properties (Figure 5). Although microplastics positively affected shoot mass, we did not find differences among shapes of the same polymer type (Figure 5, Table S4). However, these differences were evident for root mass as polyethylene (PE) and polypropylene (PP) showed statistically robust differences between shapes. That is, root mass was higher with films compared to foams made of PE and gradually increased from fibers to films and to fragments made of PP (Figure 5, Table S4). The key role of shape over polymer was also evident in soil aggregation and microbial activity. Soil aggregation was higher with fragments compared to films made of PET and with films compared to fibers made of PP, while microbial activity was lower with fragments compared to films or fibers made of PP (Figure 5, Table S4).

**Figure 5.**
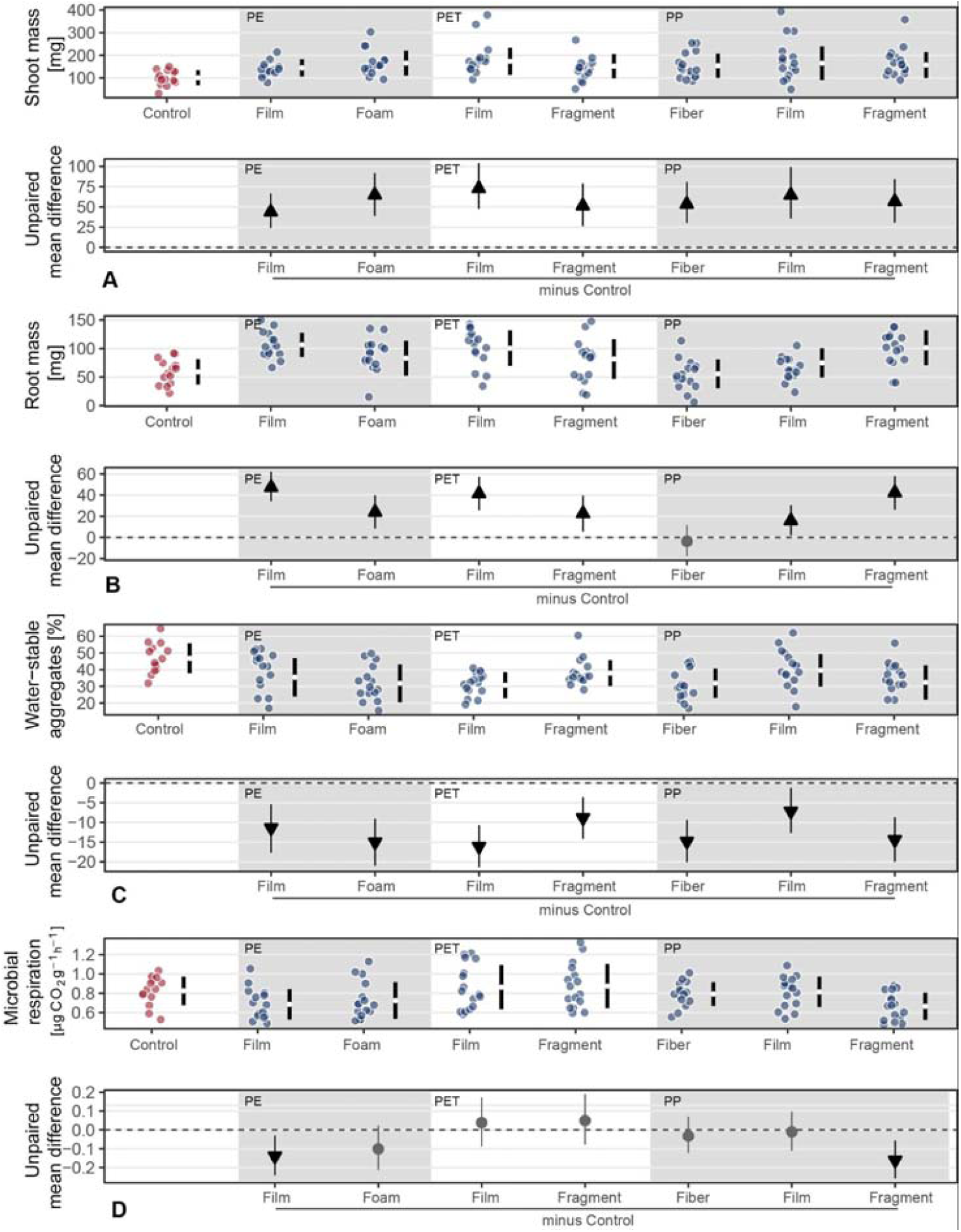
Test of the shape dissimilarity hypothesis. Effects of the same polymer type from different shapes on shoot and root masses, soil aggregation and microbial activity. Effect sizes and their variance are displayed as means and 95% confidence intervals. Effects are color-coded: gray circles indicate neutral effect sizes, black arrows with an arrow head pointing upwards or downwards indicate positive or negative effects, respectively. Horizontal dotted line indicates the mean value of the control.Polyethylene (PE), Polyethylenterephthalat (PET) and Polypropylene (PP) were selected as different shapes were made of these polymers. See statistical comparisons in Table S4; n = 7 for microplastics, n=14 for control samples.

## DISCUSSION

Our results showed that microplastic effects on plant traits and soil physical and biological properties depended on the microplastic shape, polymer type and concentration in the soil.

### Different microplastic shapes increased shoot and root biomass

Microplastics increased shoot and root masses. The detected pattern depended on microplastic shape and concentration. Overall, shoot mass increased with fiber concentration, becoming noticeable for fibers made of polypropylene (PP) and polyester (PES), while root mass increased only at highest fiber concentrations. The increase in root mass with fibers may be linked with the fact that microfibers decrease soil bulk density ^15, 32^, causing an increase in soil macroporosity and aeration ^33, 34^. This facilitates root penetration in the soil ^35^, and thus root growth. This increase in root mass could facilitate water and nutrient uptake, an effect that would be compounded with water availability, which could be increased, since microfibers enhance water holding capacity ^15^. Additionally, the increase in root mass might promote rhizodeposition and mycorrhizal associations ^36^; the latter could have contributed to the observed increase in shoot mass.

Likewise, microplastic films increased both shoot and root masses, probably due to reduction of soil bulk density ^32, 37^, and the improvement of associated soil properties ^33, 34^. Shoot and to some extent root mass were affected by microfilm concentration in a pattern opposite to that of microfibers. The decrease in shoot and root mass with microfilm concentration may be due to the creation of more channels for water movement, increasing the rate of soil evaporation ^38^. This water shortage could explain the reduction in shoot mass growth, which in our case was more evident with polypropylene (PP) and polyethylenterephthalat (PET) films. Alternatively, the increase in plant growth with microfilms could be linked with the fact that polyethylene (PE) films may promote Proteobacteria abundance ^19^, a group described as potential plant-growth promoting ^39, 40^. Similarly, shoot and root mass increased with microplastic foams and fragments as probably soil aeration and macroposity increased, which positively affected plant performance.

### Different microplastic shapes decreased soil aggregation

Microplastics of all shapes, polymer types and concentration levels decreased soil aggregation. As microplastics are incorporated into the soil matrix, they may prevent microaggregates from effectively being integrated into macroaggregates ^41^, and/or introduce fracture points into aggregates that ultimately decrease aggregate stability. Such negative effects of microfibers on aggregation has been recorded in soil with and without plants ^15, 42^. Additionally, as soil biota highly determine soil aggregation ^43^, the overall decline in this soil property may be associated with negative effects of microplastics on soil biota. Prior studies have shown that bacterial community diversity declines due to polyethylene (PE) films in soil ^20^. Indeed, Actinobacteria, which is one of the bacterial groups that most contribute to soil aggregation ^43^, reduced its abundance and richness due to the presence of microplastic films in soil ^19, 20^. Even though not addressed here, macroorganisms also contribute to soil aggregation ^44^, and may be affected by microplastics in soil. It has been observed that polyethylene (PE) and polystyrene (PS) particles can be ingested by worms ^17, 18^ and nematodes ^45^, which might affect its growth rates and caused histopathological damage, that ultimately may affected soil aggregation dynamics.

Importantly, the magnitude of microplastic effects on soil aggregation varied with concentration. There is evidence that soil aggregate stability decreases with microfiber concentration in soil without plants ^46^, however, our results show that this does not appear to be the case when introducing a plant species to the test system. We found that soil aggregation tends to increase with microfiber concentration (i.e., polypropylene (PP) and to some extent polyester (PES)), likely reflecting in part the overall positive effects on root growth, given that roots also contribute to aggregation. Similarly, foams improved soil aggregation with increasing concentration (although always lower than in soil without microplastics). By contrast, microfilms decreased soil aggregation with increasing concentration; probably since as discussed above, they can favor water loss from soils, which might have negatively affected soil aggregation.

### Microplastic shape affects soil microbial activity

Microplastic effects on microbial activity depended on microplastics shape, polymer and concentration. We expected that microbial activity would be positively correlated with soil aggregation ^44^ but we found that microplastic shape (e.g., films, foams) and its concentration modulated their relationship. Overall, as soil aggregation decreased with microplastics, the reduction in oxygen diffusion within soil pores and the effects on water flows ^47^ may explain the reduction in microbial activity as for instance, with polyethylene (PE) films at 0.4% concentration. In accordance with that, Fei et al.^20^ found that microbial activity (measured as FDAse activity) also declined with PE films addition. Likewise, the reduction of microbial activity with microplastic foams at several concentration levels, can be related with their chemical properties. Foams (e.g., polyurethane (PU) and polystyrene (PS)) are made of hazardous monomers ^8^ that could potentially affect soil biota and thus the soil microbial activity. Indeed, PS foams may contain higher concentrations of organic pollutants ^48^ not only related to its polymer structure but also related to its shape.

By contrast, we observed that polypropylene (PP) films at lower (0.1%) and high (0.4%) concentrations tended to increase microbial activity. A similar pattern was found by Liu et al. ^49^ after measuring FDAse activity in soils polluted with PP films. Previous research showed that PP fragments can release dissolved organic carbon and stimulate microbial activity ^50^. Similarly, polyester (PES) fibers at high concentrations tended to increase microbial activity. This aligns with the results of FDA activity ^15^.

### Testing the microplastic shape-related hypothesis

In this study we tested the two microplastic shape related hypotheses targeting the importance of microplastic effects caused by (i) shape mediation and (ii) shape dissimilarity. Our results support the shape mediation hypothesis that states that in addition to the shape other properties in terms of composition or additives may influence the microplastic effects ^5^. Thus, we show that equal shape of different properties had a different effect on shoot and root masses and on soil aggregation and microbial activity, as influenced by in this case, the polymer type and additives. Our results showed this for microplastic shapes such as fibers, films, foams and fragments.

Likewise, our results support the shape dissimilarity hypothesis that stated that the more dissimilar in shape from the natural population of shapes, the stronger the microplastic effects can be ^5^. Different shapes of the same polymer type (e.g., fibers, films and fragments made of polypropylene) affected the response of root mass, soil aggregation and microbial activity but not of shoot mass. The same was true for films and fragments made of Polyethylenterephthalat (PET) or films and foams made of polyethylene (PE). However, we acknowledge that to really test this hypothesis implies the use of microplastic of different shapes but identical chemical properties ^5^. Our approach deals with microplastics of different shape but only identical polymer type, as different additives (e.g., plasticizers, blowing agents, stabilizers) were likely used to obtain the desired plastic characteristics (flexibility, roughness, density, etc) ^51^ and ultimately generate the different microplastic shapes.

As microplastics may come into the soil in different shapes ^5^, polymer types ^6^ and concentrations, it is crucial to understand its effects on soil properties and plant performance, especially as the use of plastic is increasing worldwide. Our findings provide empirical evidence that in the short term, microplastics of different shapes and polymers increase shoot and root biomass, but negatively affect soil properties as aggregation and microbial activity.

However, microplastics effects on plant performance and soil properties would not only depend on the shape, polymer type and concentration levels, but also on the plant species identity. For instance, contrary to our results, Qi et al. ^52^ found that polyethylene films did not affect biomass of a wheat crop, while Lozano and Rillig ^32^ found that polyester fibers may increase biomass of some plant species while decreasing that of others in a grassland community. Likewise, contrary to our results, microfibers may rather promote soil aggregation at the plant community level ^16^. As plant species can respond differently to microplastic addition, more research is needed in order to understand the effects of shape, polymer type and concentration levels on plant performance and soil properties in a wide range of species and at the community level.

## ACKNOWLEDGMENTS

The work was funded by the German Federal Ministry of Education and research (BMBF) within the collaborative Project “Bridging in Biodiversity Science (BIBS-phase 2)” (funding number 01LC1501A). MCR additionally acknowledges funding through an ERC Advanced Grant (694368). We thank Sabine Buchert, Anja Wulf, Gabriele Erzigkeit, Jenny M. Ospina, Andrea C. Gundry and Emily Magkourilou for their assistance in the experimental setup and data collection. The authors declare no competing financial interest.

## AUTHOR CONTRIBUTIONS

YML and MCR conceived the ideas and designed methodology; YML, TL established and maintained the experiment in the greenhouse; LTL analyzed the soil aggregation. AL designed the figures. YML analyzed the data and wrote the first draft of this manuscript. All authors contributed to the final version and gave final approval for publication.

## SUPPORTING INFORMATION

**Table S1.** Microplastic shape effect on shoot and root masses, soil aggregation and microbial activity. Results of linear model, and multiple comparisons by using the Dunnett test. Values in bold denote a significant effect (p<0.05) of the treatment on the dependent variable.

| Linear model | Shoot mass |  |  | Root mass |  | Soil aggregation |  | Microbial activity |  |
| --- | --- | --- | --- | --- | --- | --- | --- | --- | --- |
|  | df | F value | p value | F value | p value | F value | p value | F value | p value |
| Treatment | 4 | 10.62 | <0.001 | 21.99 | <0.001 | 8.197 | <0.001 | 0.949 | 0.436 |

| Multiple comparisons (Dunnett) | Shoot mass |  | Root mass |  | Soil aggregation |  | Microbial activity |  |
| --- | --- | --- | --- | --- | --- | --- | --- | --- |
| Treatment - control >= 0 | z value | p-value | z value | p-value | z value | p-value | z value | p-value |
| Fibers | 2.676 | 0.011 | 0.551 | 0.513 | -5.313 | <0.001 | -0.415 | 0.970 |
| Films | 5.486 | <0.001 | 5.183 | <0.001 | -4.367 | <0.001 | -0.768 | 0.799 |
| Foams | 4.325 | <0.001 | 5.066 | <0.001 | -3.583 | <0.001 | -1.698 | 0.221 |
| Fragments | 5.037 | <0.001 | 4.320 | <0.001 | -4.811 | <0.001 | -1.047 | 0.599 |

**Table S2.**
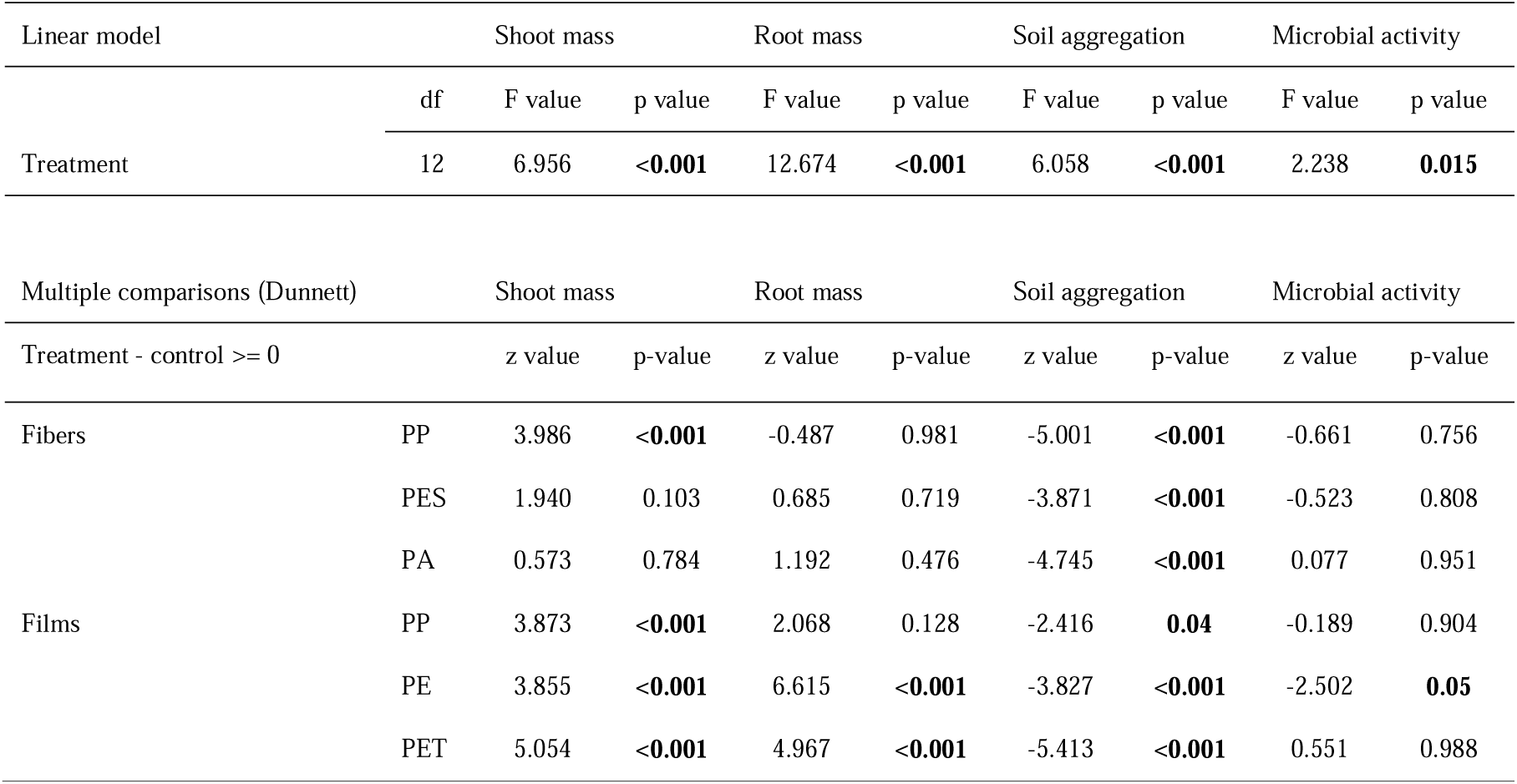
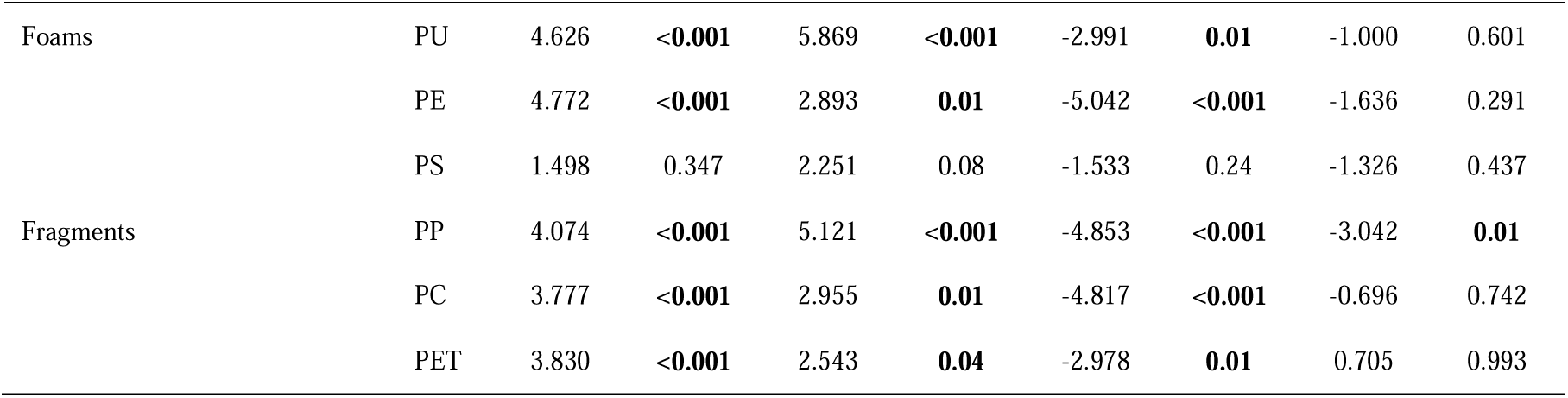
Polymer type effect on shoot and root masses, soil aggregation and microbial activity. Results of linear model, and multiple comparisons by using the Dunnett test. Polypropylene (PP); Polyester (PES); Polyamide (PA); Polyethylene (PE); Polyethylenterephthalat (PET); Polyurethane (PU); Polystyrene (PS); Polycarbonate (PC). Values in bold denote a significant effect (p<0.05) of the treatment on the dependent variable.

| Linear model |  | Shoot mass |  | Root mass |  | Soil aggregation |  | Microbial activity |  |
| --- | --- | --- | --- | --- | --- | --- | --- | --- | --- |
|  | df | F value | p value | F value | p value | F value | p value | F value | p value |
| Treatment | 12 | 6.956 | <0.001 | 12.674 | <0.001 | 6.058 | <0.001 | 2.238 | 0.015 |

| Multiple comparisons (Dunnett) |  | Shoot mass |  | Root mass |  | Soil aggregation |  | Microbial activity |  |
| --- | --- | --- | --- | --- | --- | --- | --- | --- | --- |
| Treatment - control >= 0 |  | z value | p-value | z value | p-value | z value | p-value | z value | p-value |
| Fibers | PP | 3.986 | <0.001 | -0.487 | 0.981 | -5.001 | <0.001 | -0.661 | 0.756 |
|  | PES | 1.940 | 0.103 | 0.685 | 0.719 | -3.871 | <0.001 | -0.523 | 0.808 |
|  | PA | 0.573 | 0.784 | 1.192 | 0.476 | -4.745 | <0.001 | 0.077 | 0.951 |
| Films | PP | 3.873 | <0.001 | 2.068 | 0.128 | -2.416 | 0.04 | -0.189 | 0.904 |
|  | PE | 3.855 | <0.001 | 6.615 | <0.001 | -3.827 | <0.001 | -2.502 | 0.05 |
|  | PET | 5.054 | <0.001 | 4.967 | <0.001 | -5.413 | <0.001 | 0.551 | 0.988 |
| Foams | PU | 4.626 | **<0.001** | 5.869 | **<0.001** | -2.991 | **0.01** | -1.000 | 0.601 |
|  | PE | 4.772 | **<0.001** | 2.893 | **0.01** | -5.042 | **<0.001** | -1.636 | 0.291 |
|  | PS | 1.498 | 0.347 | 2.251 | 0.08 | -1.533 | 0.24 | -1.326 | 0.437 |
| Fragments | PP | 4.074 | **<0.001** | 5.121 | **<0.001** | -4.853 | **<0.001** | -3.042 | **0.01** |
|  | PC | 3.777 | **<0.001** | 2.955 | **0.01** | -4.817 | **<0.001** | -0.696 | 0.742 |
|  | PET | 3.830 | **<0.001** | 2.543 | **0.04** | -2.978 | **0.01** | 0.705 | 0.993 |

**Table S3.**
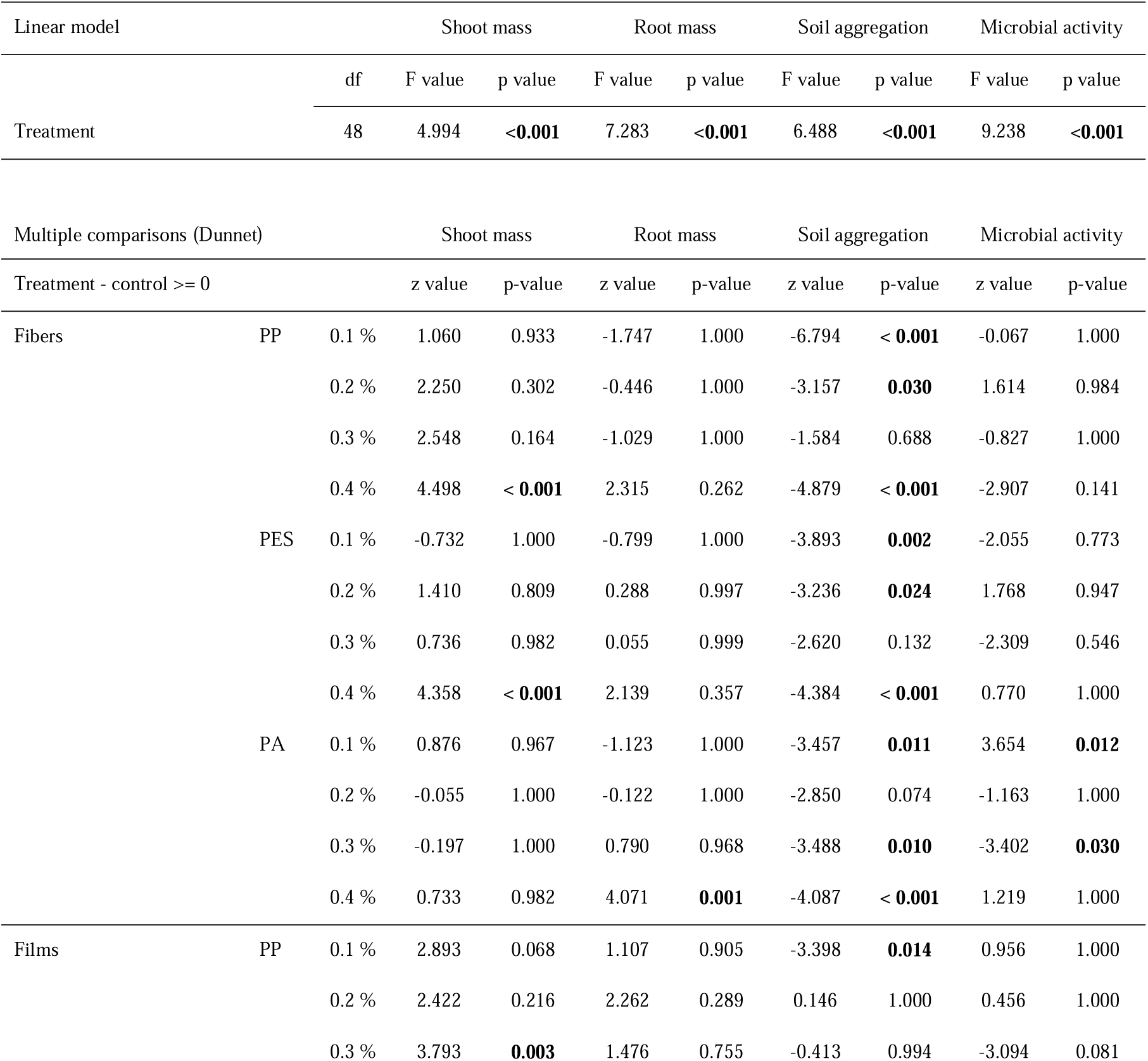
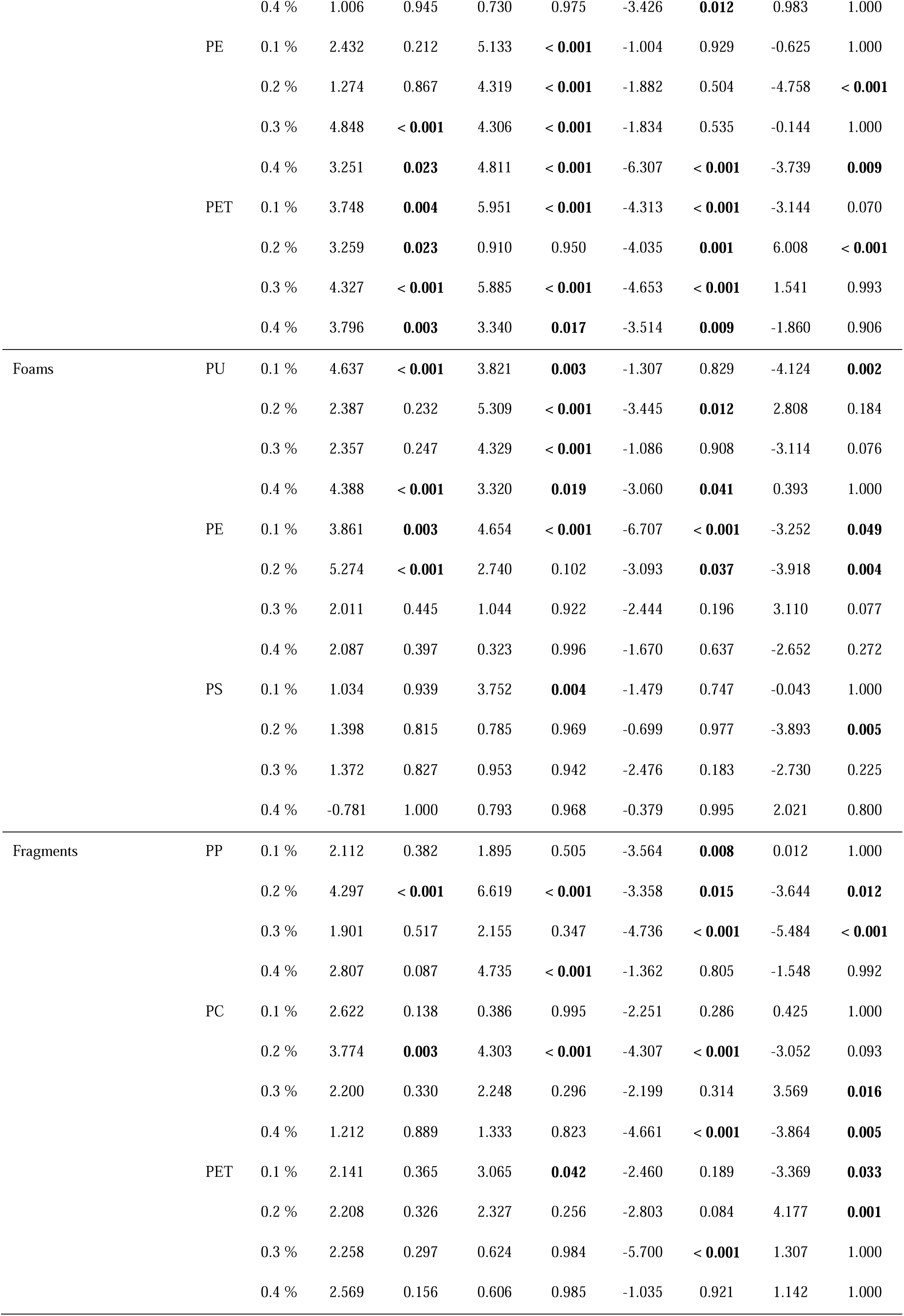
Microplastic concentration effect on shoot and root masses, soil aggregation and microbial activity. Results of linear model, and multiple comparisons by using the Dunnett test. Polypropylene (PP); Polyester (PES); Polyamide (PA); Polyethylene (PE); Polyethylenterephthalat (PET); Polyurethane (PU); Polystyrene (PS); Polycarbonate (PC). Concentration levels: 0.1% to 0.4%. Values in bold denote a significant effect (p<0.05) of the treatment on the dependent variable.

**Table S4.** Test of the shape dissimilarity hypothesis. Effects of the same polymer type from different shapes on shoot and root masses, soil aggregation and microbial activity. Results of analysis of variance (aov), and multiple comparisons by using the Tukey test. Polyethylene (PE), Polyethylenterephthalat (PET) and Polypropylene (PP) were selected as different shapes were made of these polymers. Values in bold denote a significant effect (p<0.05) of the treatment on the dependent variable.

| Anova (aov) |  | Shoot mass |  | Root mass |  | Soil aggregation |  | Microbial activity |  |
| --- | --- | --- | --- | --- | --- | --- | --- | --- | --- |
|  | df | F value | p value | F value | p value | F value | p value | F value | p value |
| Treatment | 6 | 0.802 | 0.57 | 10.97 | **<0.001** | 3.538 | **0.002** | 3.499 | **0.003** |

| Multiple comparisons (Tukey) |  | Shoot mass |  | Root mass |  | Soil aggregation |  | Microbial activity |  |
| --- | --- | --- | --- | --- | --- | --- | --- | --- | --- |
| Treatment |  | t value | p-value | t value | p-value | t value | p-value | t value | p-value |
| PE | Films vs Foams | 1.437 | 0.777 | -3.384 | **0.014** | -1.208 | 0.888 | 0.643 | 0.994 |
| PET | Films vs Fragments | -1.707 | 0.607 | -2.183 | 0.305 | 3.549 | **0.008** | 0.137 | 1.000 |
| PP | Fibers vs Films | 0.330 | 1.000 | 2.912 | **0.054** | 3.164 | **0.029** | 0.464 | 0.999 |
|  | Fibers vs Fragments | 0.264 | 1.000 | 6.245 | **<0.001** | 0.175 | 1.000 | -2.836 | 0.075 |
|  | Films vs Fragments | -0.107 | 1.000 | 3.571 | **0.007** | -2.758 | 0.088 | -2.935 | **0.058** |

